# 3D DNA structural barcode copying and random access

**DOI:** 10.1101/2020.11.27.401596

**Authors:** Filip Bošković, Alexander Ohmann, Ulrich F. Keyser, Kaikai Chen

## Abstract

Three-dimensional (3D) DNA nanostructures built via DNA self-assembly have established recent applications in multiplexed biosensing and storing digital information. However, a key challenge is that 3D DNA structures are not easily copied which is of vital importance for their large-scale production and for access to desired molecules by target-specific amplification. Here, we build 3D DNA structural barcodes and demonstrate the copying and random access of the barcodes from a library of molecules using a modified polymerase chain reaction (PCR). The 3D barcodes were assembled by annealing a single-stranded DNA scaffold with complementary short oligonucleotides containing 3D protrusions at defined locations. DNA nicks in these structures are ligated to facilitate barcode copying using PCR. To randomly access a target from a library of barcodes, we employ a non-complementary end in the DNA construct that serves as a barcode-specific primer template. Readout of the 3D DNA structural barcodes was performed with nanopore measurements. Our study provides a roadmap for convenient production of large quantities of self-assembled 3D DNA nanostructures. In addition, this strategy offers access to specific targets, a crucial capability for multiplexed single-molecule sensing and for DNA data storage.

## 1. Introduction

DNA nanotechnology offers structural control with sub-nanometer precision to build designed three-dimensional (3D) nanostructures by DNA self-assembly relying on Watson-Crick base pairing.^[1]^ This striking advantage was exploited in numerous ways to create different functional DNA nanostructures.^[1–3]^ One useful aspect that DNA nanotechnology offers is to precisely position 3D DNA nanostructures along a DNA strand in a programmable arrangement.^[2,4]^ By annealing a several kilobase-long, single-stranded (ssDNA) ‘scaffold’ strand with 38-48 nt complementary oligonucleotides containing DNA nanostructures, uniquely addressable 3D DNA structural barcodes can be created.^[2]^ It is comparable to the DNA sequence barcode which has been employed to specifically label DNA fragments for DNA sequencing and genomic library preparation.^[5,6]^ However, with the advance of DNA nanotechnology, 3D DNA structural barcodes emerged as a potential alternative by encoding information in the structures rather than in the sequences. They are more robust in carrying embedded information as multiple single-nucleotide changes could cause misinterpretation of barcode DNA sequence,^[7]^ whereas the barcode readout of the 3D structure will not be affected. Furthermore, it is easier and more accurate to read the codes on the 3D DNA structures because they can be designed with more significant difference in size compared to the small difference in single nucleotides. Some of the versatile applications of 3D DNA structural barcodes are data storage in DNA nanostructures^[8]^ and multiplexed sensing of single proteins both *in vitro*^[2]^ and *in vivo*.^[9]^ DNA barcodes have been shown to be promising tools for multiplex intracellular protein sensing.^[9]^ Combined with fluorescent labelling, they have additionally shown tremendous applications in the field of DNA-based point accumulation for imaging in nanoscale topography (DNA-PAINT) to create a fluorescent barcode.^[10,11]^ 3D DNA structural barcodes have also a great potential to be used in whole-genome mapping strategies, such as single-molecule optical mapping.^[12]^

To read out the information stored in 3D DNA structural barcodes, nanopore-based single-molecule sensing has proven to be a powerful tool for numerous applications.^[2,8,13]^ The principle of nanopore sensing is based on passing a single molecule through a small confinement by applying a voltage across it. The translocation of the charged molecule driven by the electric field creates a current blockage which is proportional to the molecule’s shape, volume, and charge allowing us to read out structural information in a label-free manner. Recently, we have pushed this even further by using DNA nanostructures for an innovative data storage approach as an alternative to data storage in DNA sequence^[8]^ including the ability to rewrite and securely store data in so-called DNA hard drives.^[13]^

3D DNA structural barcodes are not easy to be copied because of their complexity^[14]^ compared to a DNA sequence.^[15]^ While 3D DNA structural barcodes may be copied via transformation of bacteria with the self-replicating unit,^[16]^ an approach based on polymerase chain reaction (PCR) amplification is more valuable as it is a standardized laboratory method.^[17]^ However, a key challenge is the asymmetry of the 3D DNA structural barcode design. While the scaffold strand is linear, its complement contains individual oligonucleotides where some of them include 3D DNA protrusions. A 3D DNA structural barcode composed of multiple, closely spaced DNA nanostructures will contain repetitive DNA.^[14]^

In addition, accessing a desired 3D DNA nanostructure barcode in a complex mixture of similar barcodes is still a huge challenge because of their similar design and usage of the same DNA scaffold. Random access is of paramount significance for targeting and enrichment of the 3D DNA structural barcode of interest, for example, to read only desired data files and not the whole archive of stored data. As single-molecule methods typically require longer measurement times or multiple parallel measurements, significant amounts of library material could be wasted to detect one file. Hence, random access is critical for more realistic applications of 3D DNA structural barcodes carrying digital information.

To overcome those emerging challenges, we successfully developed barcode copying and random access of single barcodes from the library of 3D DNA structural barcodes in this study. Firstly, we designed and assembled a library of 3D DNA structural barcodes where a 3D bit is composed of multiple dumbbell-shaped, double-hairpin DNA protrusions (see **Figure 1A** and Supplementary Figures S1 and S2 for details). The design of the barcodes was verified with nanopore sensing and the expected barcode lengths were confirmed with agarose gel electrophoresis. Subsequently, we successfully copied a barcode with six 3D bits with exponential PCR amplification. Finally, we designed a library of barcodes with an end-specific primer that we employed to access desired barcode by barcode-specific linear PCR amplification and verifying the output with nanopore sensing. Our study introduces an easier and more affordable approach towards fabricating data storage archives and barcode libraries for various biomolecule sensing purposes using 3D DNA structural barcodes.

**Figure 1.**
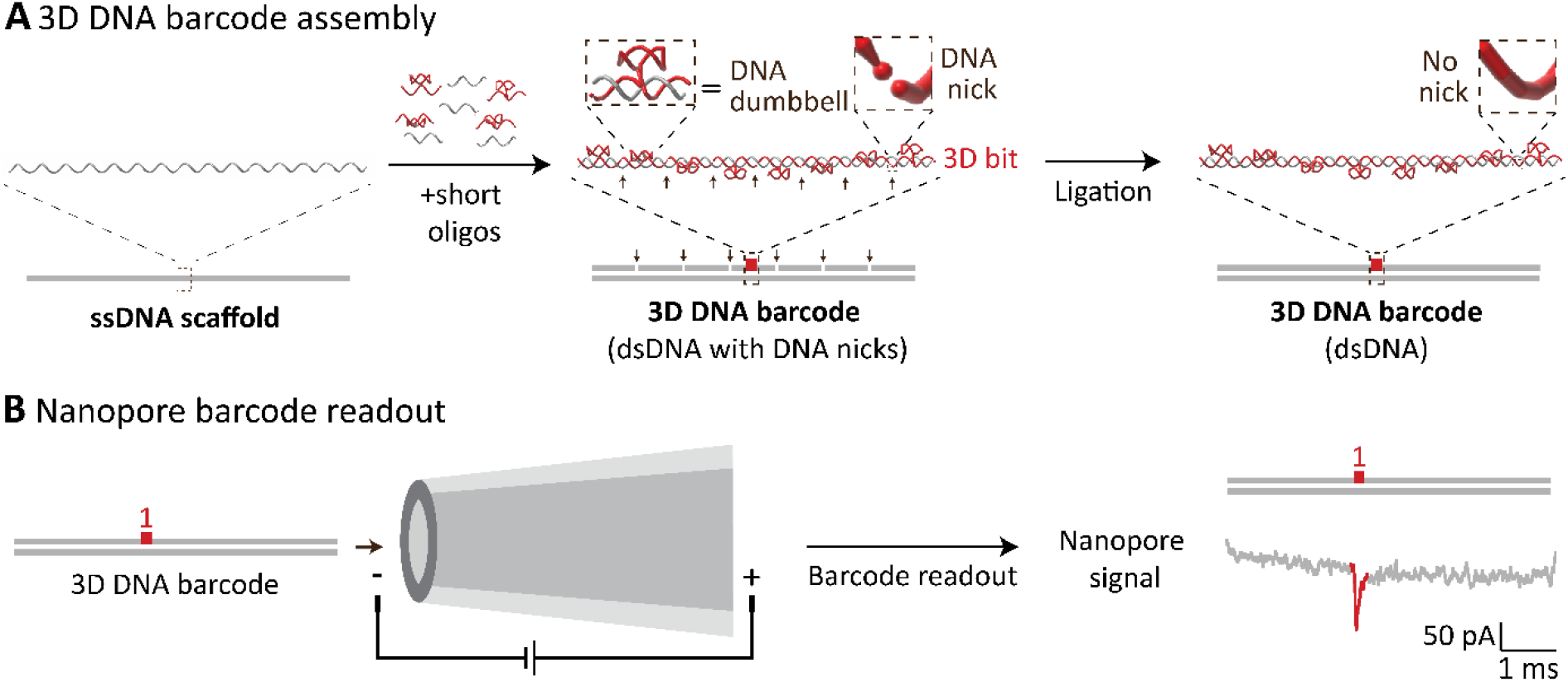
3D DNA structural barcode assembly and nanopore readout. **A** DNA barcodes are assembled by hybridizing single-stranded DNA scaffolds (7228 nt) with short complementary oligonucleotides (38-48 nt). Specific oligonucleotides are designed to form double DNA hairpins (DNA dumbbells). Each 3D bit of the DNA barcode consists of eight, closely spaced DNA dumbbells. DNA nicks (black arrows), resulting from separate oligos hybridized to the scaffold, are ligated using a T4 DNA ligase to form the final double-stranded 3D DNA barcode. **B** Negatively charged 3D DNA barcodes pass through the nanopore creating a barcode-specific nanopore signal as the readout. The 3D bit is detected as a downward peak in the current assigned to be ‘1’.

## 2. Results

### 2.1. 3D DNA structural barcode design, assembly and readout

To create 3D DNA barcodes,, we first prepared single-stranded DNA scaffold (7228 nt) by linearizing circular M13mp18 DNA and subsequently annealing this linear scaffold (Figure 1A left) with short, 5’ phosphorylated oligonucleotides (sequence of oligonucleotides are listed in Supplementary Table S1). The particular barcode was then assembled by designing selected complementary oligonucleotides witha double hairpin structure (Figure 1A middle) termed ‘DNA dumbbell’ (sequences listed in Supplementary Table S2-5). This DNA nanostructure is formed of two hairpins each consisting of 5 bp and 4 dT loop as shown in Supplementary Figure S2. A single 3D bit is created by closely spacing eight DNA dumbbell nanostructures together along the long DNA scaffold (Supplementary Figure S1). After annealing with the short oligonucleotides, we obtained a double-stranded 3D DNA barcode with multiple DNA nicks (indicated with vertical black arrows in Figure 1A middle). We copied the structure using PCR by subsequently linking the separate oligonucleotides together by ligating the DNA nicks using T4 DNA ligase. The ligation forms a long continuous dsDNA strand with designed 3D bits protruding from it (Figure 1A right).

The correct assembly of 3D DNA structural barcodes was verified by reading out the barcode using nanopore sensing (Figure 1B). The negatively charged 3D DNA barcodes were electrophoretically driven through the nanopore by the applied electric field thereby causing a measurable, barcode-specific modulation of the ionic current. The nanopore sensing method can detect protrusions along the DNA as an additional, distinct downward peak in current signal as shown in Figure 1B (right). The position of the 3D bit in the design then corresponds to the peak position in the ionic current recording due to a relatively constant DNA translocation velocity.^[18]^ The presence of a 3D bit, indicated by a downward peak, is assigned as ‘1’, and the absence of a peak is assigned as a 3D bit ‘0’.

### 2.2. Nanopore microscope verification of designed 3D barcode library

Before copying, we first verified each barcode design individually using nanopore sensing. For this, we collected datasets for each barcode design, isolated single barcode translocations along the current trace, and used only unfolded barcode translocations for further analysis (for detailed analysis workflow see Supplementary Information Section 3. We prepared four barcode designs: one barcode with 6-bits and another three with each 3-bits (**Figure 2**). Figure 2A illustrates the 6-bit DNA barcode design together with an example of the typical, corresponding nanopore signal. Each 3D bit is highlighted in red and corresponds to a downward peak in the nanopore recording. On approximately 93 % of events, six 3D bits corresponding to our designed barcode could be determined (Figure 1A), demonstrating the compelling accuracy of our approach.

**Figure 2.**
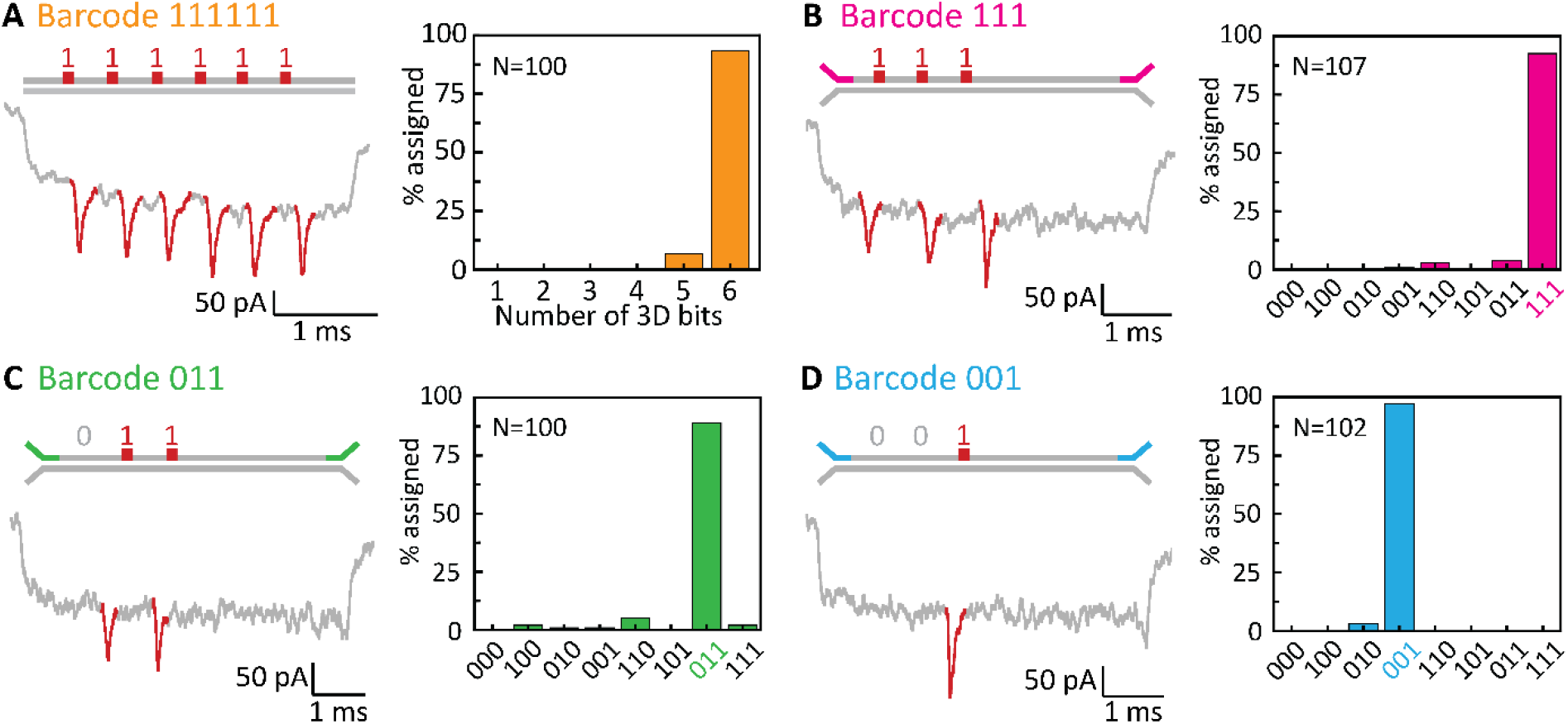
3D structural barcode characterisation. We tested four different barcode signatures represented by a varying number of 3D bits (red). **A** Barcode ‘111111’ contains six equally spaced 3D bits. Exemplary nanopore readout showing six peaks in the current signal is depicted below. 93 % of counted events were correctly assigned with six 3D bits. **B-D** Three different barcodes containing three (B), two (C) and one (D) 3D bit were verified with nanopore barcode readout yielding 93 %, 90 %, and 97 % accuracy, respectively. Exemplary events are depicted below each design. Non-complementary terminal sequences offer barcode-specific amplification using strand-specific primers (discussed in Figure 4).

For the 3-bit systems, we designed three 3D DNA structural barcodes encoding for ‘111’, ‘011’ and ‘001’, by having three, two or one 3D units respectively. Illustrations of barcodes ‘111’, ‘011’ and ‘001’ are shown in Figures 2B-D respectively, with representative events and the statistical analysis of the assigned readout to a certain barcode. Additional examples are shown in Supplementary Figure S4 with corresponding event statistics in Supplementary Figure S8. All three designs demonstrated a readout with > 90 % correct readout, further establishing the robustness of our method. An additional feature of 3-bit barcodes is the non-complementary ends (Supplementary Table S6) serving as a template for barcode-specific copying and access which will be discussed later in this study.

### 2.3. Copying and quantification of 3D DNA structural barcodes

We demonstrate the copying capability by amplifying the barcode ‘111111’ with six 3D bits introduced in Figure 2A using LongAmp® Taq DNA Polymerase (NEB). Using a thermocycler for the PCR reaction, the double-stranded DNA barcode was denatured by heat to obtain two ssDNA strands. These were subsequently annealed with forward and reverse primers (for sequences see Supplementary Table S3) followed by elongation step of the primer strands. Before copying, the majority of events contain six 3D units and the barcode consists of two strands B (containing DNA dumbbells) and S (DNA scaffold), as verified in Figure 2A. In the first amplification cycle, strands S1 and S2 are copied to obtain the dsDNA constructs S1S1’ and S2S2’ (S1’ and S2’ are the complementary strands of S1 and S2, respectively) using the forward and the reverse primers. After the additional 29 cycles of exponential PCR copying of initial asymmetric DNA strands, four different strand combinations could exist. S1S2 and S2’S1’ should have identical ionic current nanopore traces with the correct reading of ‘111111’ while the S2S2’ combination consists of complementary DNA strands without the 3D bits (**Figure 3A**). The last combination is S1S1’ that might form dsDNA constructs with or without protrusions due to the full complementary of protrusions.

**Figure 3.**
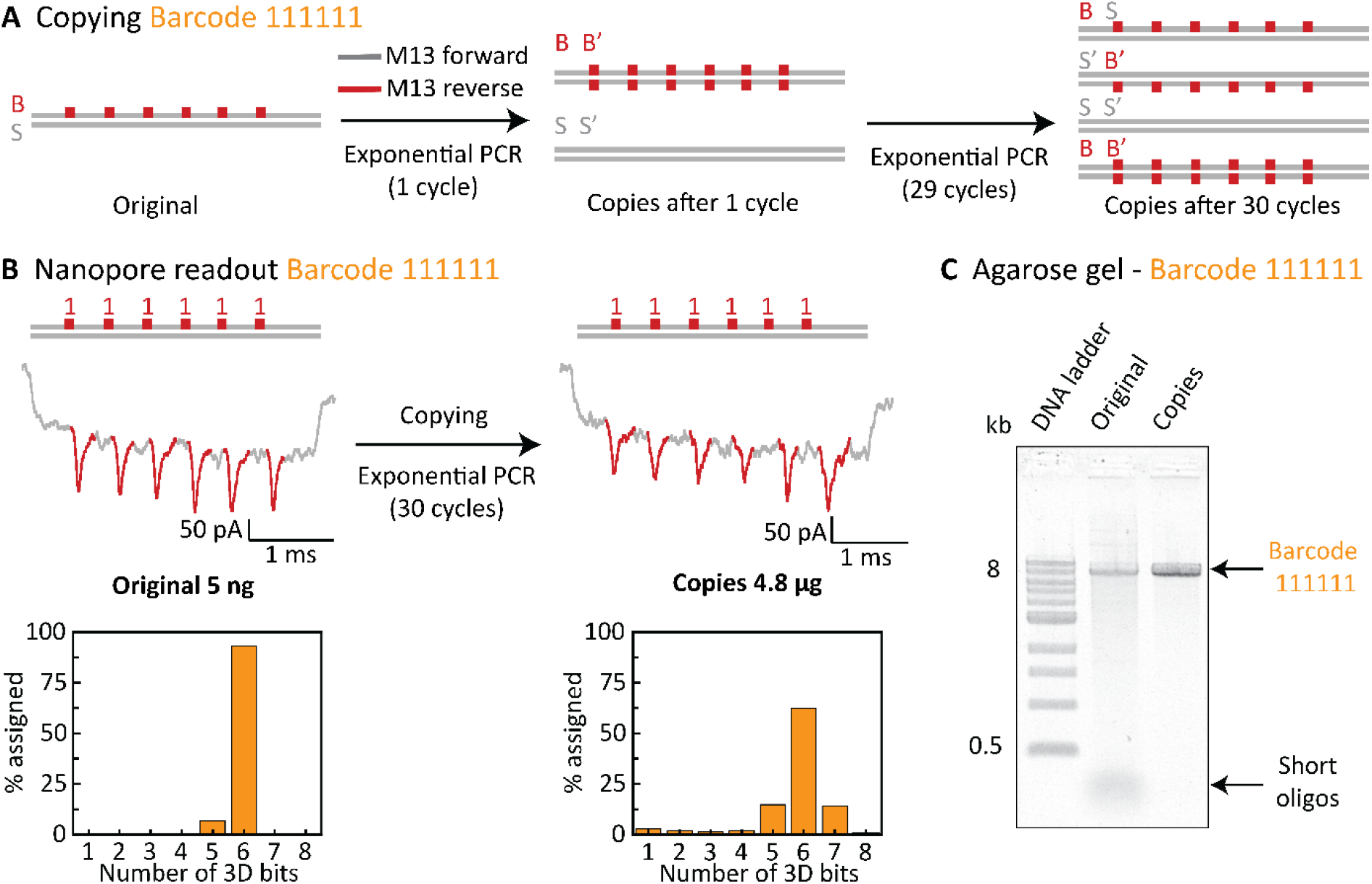
Copying of the 3D DNA barcodes using PCR amplification. **A** Barcode ‘111111’ is comprised of strands B (containing 3D bits) and S (DNA scaffold). Using a pair of primers (forward and reverse primers), B and S are initially replicated to their complements B’ and S’. Subsequent exponential PCR amplification primarily yields structurally comparable barcodes as well as constructs with complementary strands. **B** PCR amplification increased the number of copies of barcode ‘111111’ by almost 1,000-fold increase almost three orders of magnitude. Importantly, the nanopore readout of the barcode ‘111111’ copies remained comparable to its original, as illustrated by exemplary nanopore current traces. **C** Agarose gel electrophoresis verifies the consistent length of 3D DNA barcode ‘111111’ before and after copying at ∼8 kb, as expected.

3D DNA barcodes were verified by nanopore sensing before and after copying confirming that the six 3D-bit barcode features were still preserved after PCR copying (Figure 3B). Importantly, we were able to amplify the initial 5 ng of original barcodes by close to 1,000-fold, yielding 4.8 μg of copied DNA barcode structures. Statistical nanopore single-molecule readouts confirm that the majority (> 60 %) of counted events are still correctly assigned. Additional examples of the correct barcode readout after amplification are shown in Supplementary Figure S6.

To support the single-molecule results of the nanopore experiments, we additionally performed agarose gel electrophoresis as a bulk analysis method to verify the DNA barcode lengths before and after copying. By comparing the observed gel bands to a DNA ladder, we could confirm that the DNA barcodes before and after amplification agree with the expected length of close to 8 kb (Figure 3C), thus furthermore demonstrating the successful copying process.

### 2.4. Random access of a desired barcode from a barcode library

As introduced above, the 3-bit barcode structures were designed to contain a non-complementary, unique end sequence which allows us to use a barcode-specific primer to access a single barcode from the library of mixed barcodes (**Figure 4A**). We demonstrate the access of a desired target barcode from a library by amplifying one barcode from an equimolar mixture of three 3-bit barcodes. For this, barcodes ‘111’, ‘011’, and ‘001’ were mixed in equimolar concentrations. To perform the linear PCR amplification, we added barcode ‘111’-specific primer strands and the “M13mp18 forward primer”. After performing 60 amplification cycles, the output of this reaction should be mainly composed of the amplified barcode ‘111’.

**Figure 4.**
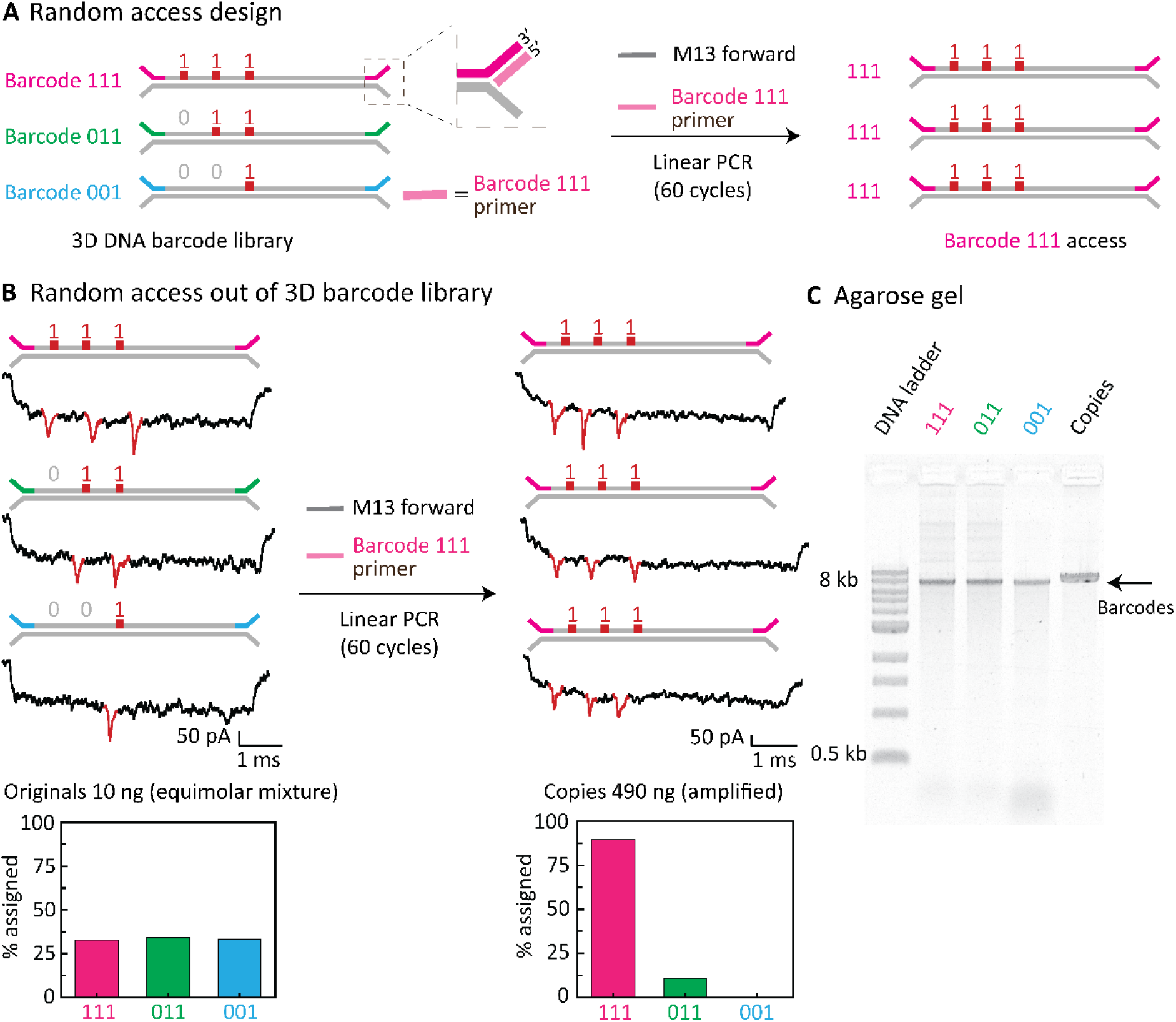
Random access to a specific 3D DNA barcode by selective copying. **A** 3D DNA barcode library represents the equimolar mixture of barcodes ‘111’, ‘011’, and ‘001’. Barcode-specific non-complementary ends are highlighted in different colours. Random access is achieved by selective PCR copying using a barcode-specific pair of primers. As an example, barcode ‘111’ is amplified for 60 cycles, with DNA amount indicated in nanograms. **B** 3D DNA barcode library including exemplary current traces before and after copying highlighting the increased nanopore readout of barcode ‘111’ after its selective amplification from an equimolar barcode mixture. **C** Agarose gel electrophoresis verifies the size of each barcode design before copying as well as in the purified and diluted mixture after PCR. 1 kb ladder indicates that barcodes are close to 8 kb.

We validated this by nanopore measurements before and after copying of barcode ‘111’ from the described barcode library (Figure 4B; for example events see Supplementary Figures S5 and S7). Before amplification all barcodes are read out with an equal probability (∼30 %), as expected from an equimolar barcode mixture. However, after amplification the vast majority (> 88 %) of barcodes are assigned to the desired amplified barcode ‘111’. Three representative events before and after amplification are shown in Figure 4B. All three different barcodes are detected before the amplification but after selective copying, barcode ‘111’ is in excess which is quantitatively estimated from Nanodrop spectrophotometer (Figure 4B). Furthermore, we employed agarose gel electrophoresis again to confirm that the barcodes display the expected band at ∼8 kb before and after selective amplification (Figure 4C).

## 3. Discussion

In this study, we demonstrated that our 3D DNA structural barcode designs can be successfully copied which we verified by single-molecule nanopore analysis and bulk agarose gel electrophoresis analysis. Secondly, we demonstrated that by employing barcode-specific primers, we can randomly access a specific target barcode from a library of constructs.

We successfully copied the six-bit 3D DNA structural barcode using exponential PCR that should create an equimolar mixture of four DNA strands that can hybridize into multiple copies of the original barcode ‘111111’. Still, the complexity of the PCR reaction can lead to a different number of copied DNA dumbbells (= 3D bits) due to the repetitive nature of the sequence forming DNA protrusions.^[19]^ However, we show that our approach is robust and that the barcode readout is correct regardless imperfections of PCR amplification.

A DNA protrusion can be replaced with a non-complementary DNA loop that contains a binding site to which additional DNA nanostructures^[20]^ or biotinylated strands.^[13]^ Additionally, the binding site can be in the loop region of DNA stem-loop structure and both structures could be copied and accessed in a similar way as DNA dumbbells and serve as additional ways to create 3D DNA structural barcodes. As it has been shown previously one can rewrite such barcode that further can be used as a unique on-demand programmable platform for barcoding.^[13]^

Self-assembled structural 3D DNA barcodes are one of the most promising choices to create durable barcodes for nanopore sensing as they have several advantages. The barcode readout in nanopores should not be affected if several DNA dumbbells are missing.^[2]^ DNA nanostructures are intrinsically unaffected by a few nucleotide substitutions in the sequence of the barcode because this would not cause a significant difference in the barcode structure. Using DNA self-assembly or strand displacement reaction one can easily change the position and number of DNA structures in a barcode, as we previously demonstrated.^[2,13]^

Random access in DNA data libraries is of particular interest for DNA data storage.^[21]^ Our work will pave the way for data storage in DNA nanostructures, a new approach shown recently as an alternative to DNA sequence-based data storage.^[13,22–24]^ The same system can be employed to access a stored data file of interest by copying it. This approach can offer large quantities of barcodes for the product manufacturing monitoring that combined with nanopore sensing offers fast barcode retrieval.^[13,25]^ We created multiple copies of a specific data file written in a 3D DNA structural annotation that can expand the potential of DNA nanostructures and nanopores as a feasible way to archive and access data.

Our study will propel the field towards sustainable, larger-scale application of 3D structural barcodes for multiplexed single-molecule sensing of biomolecules as has been introduced previously.^[2]^ Production of larger quantities of such barcodes has been one of limiting factors for industrial-scale applications of self-assembled DNA systems.^[26,27]^

## 4. Experimental Section/Methods

### 3D DNA barcode assembly

We fabricate the barcodes by annealing a long linear ssDNA scaffold with short complementary oligonucleotides (Integrated DNA Technologies, Inc.) of which some have a fully complementary sequence (Supplementary Table S1). A select number of these oligos additionally contain DNA protrusions at specific positions (Figure 1A, Supplementary Tables S2-S5). The scaffold linerization protocol was previously described.^[2]^ The oligonucleotide set for assembling a specific barcode design was prepared by mixing the required oligonucleotides (200 nM final concentration per oligonucleotide) and phosphorylating the 5’ ends of the oligonucleotides using the T4 polynucleotide kinase (PNK) kit (New England Biolabs (NEB)). For phosphorylation of oligonucleotides 50 μL of 2000 picomoles of oligonucleotides were mixed with 10 μL 10× ligase buffer, 10 μL 10 mM ATP, 7 μL T4 PNK and 23 μL of nuclease-free water. The mix was incubated overnight at 37 °C and heat inactivated at 65 °C for 20 minutes.

The assembly mix for 3D DNA structural barcodes was composed of: 8 μL linearized M13mp18 DNA scaffold (100 nM); 40 μL phosphorylated oligonucleotide mix (100 nM); 6 μL 100 mM MgCl_2_; 1.2 μL 100 mM Tris-HCl (pH 8.0), 10 mM EDTA; 4.8 μL Mili-Q water. The mixture was then heated to 70 °C followed by a linear cooling ramp to 25 °C over 50 minutes.

Immediately after the assembly, annealed barcodes were purified from excess oligonucleotides via spin filtration using100 kDa Amicon filters by mixing the annealed mixture and adding washing buffer (10 mM Tris-HCl pH 8.0, 0.5 mM MgCl_2_) up to 500 μL and centrifugated at 9,000×G for 10 minutes. This step was repeated twice. The purified 3D DNA structural barcode was retrieved by reversing the filter and centrifugation for 2 minutes at 1,000×G.

Finally, DNA nicks left between hybridized oligonucleotides was ligated using the T4 DNA ligation kit (NEB) by mixing 100 ng of purified barcodes with 2 μL of 10× ligase buffer, 2 μL of T4 DNA ligase and filled up to 20 μL with nuclease-free water. The ligation reaction was incubated overnight at 16 °C, following by inactivation at 65 °C for 10 minutes. The product of the ligation reaction was purified using the Monarch® PCR & DNA Clean up Kit (5 μg), according to the manufacturer’s instructions.

### Copying of 3D DNA structural barcodes with exponential amplification

To demonstrate the exponential barcode PCR amplification, we employed a barcode design with six bits (‘111111’). Exponential PCR amplification was performed using the LongAmp Taq PCR kit (NEB; for primer sequences see Supplementary Table S7) at an initial DNA amount of ∼5 ng per reaction as estimated using a NanoDrop™ spectrophotometer. Samples were initially denatured at 94 °C for 4 min and subsequently amplified in 30 cycles of 94 °C (30 s), 54 °C (30 s), and 72 °C (7.5 min; ∼50 s/kb). The elongation time is adapted to the length of the strand containing DNA dumbbells according to a rate of ∼1 kb per second. After the amplification cycles a final extension of 72 °C for 10 min was performed, followed by storage at 4 °C. Samples were then purified using the Monarch® PCR & DNA Cleanup Kit (5 μg).

### Random access to target 3D DNA structural barcode with linear amplification

To demonstrate the exponential barcode PCR amplification, we employed a barcode design with six bits (‘111111’). Exponential PCR amplification was performed using the LongAmp Taq PCR kit (NEB; for primer sequences see Supplementary Table S7) at an initial DNA amount of ∼5 ng per reaction as estimated using a NanoDrop™ spectrophotometer. Samples were initially denatured at 94 °C for 4 min and subsequently amplified in 30 cycles of 94 °C (30 s), 54 °C (30 s), and 72 °C (7.5 min; ∼50 s/kb). The elongation time is adapted to the length of the strand containing DNA dumbbells according to a rate of ∼1 kb per second. After the amplification cycles a final extension of 72 °C for 10 min was performed, followed by storage at 4 °C. Samples were then purified using the Monarch® PCR & DNA Cleanup Kit (5 μg).

### Agarose gel electrophoresis

Lengths of the barcode constructs were verified using 0.8 % (w/v) agarose (Sigma-Aldrich, BioReagent) gel electrophoresis in a 0.5× Tris–borate–EDTA (TBE, pH 8.0) buffer. 150 ng sample were loaded using a 6×SDS-free, purple loading dye (NEB). The gel was run for 90 minutes at 4°C at 60V, post-stained in 3×GelRed (Biotium), and imaged using a GelDoc-It™ UV imaging system. A 1kb ladder (NEB) was used in all experiments. The grayscale was inverted in the ImageJ software and the background was subtracted using the integrated rolling ball method at a radius of 150 pixels.

### Nanopore measurement and data analysis

To detect the designed barcodes and their bits, we used 14 ± 3 nm glass nanopores fabricated by pulling quartz capillaries with filaments (0.5 mm outer diameter and 0.2 inner diameter, Sutter Instrument, USA) using a laser-assisted puller (P-2000, Sutter Instruments).^[2]^

The details of nanopore measurement, nanopore setup and data analysis were used as employed previously (Supplementary Figure S3).^[2]^

## Acknowledgements

We thank Pablo A. Vargas Rosales for critical reading of the manuscript. Funding: U.F.K., K.C. acknowledge funding from an European Research Council (ERC) consolidator grant (DesignerPores No. 647144). F.B. acknowledges funding from George and Lilian Schiff Foundation Studentship, the Winton Programme for the Physics of Sustainability PhD Scholarship and St John’s College Benefactors’ Scholarship. A.O. acknowledges funding from Cambridge Trust Vice Chancellor’s Award.

