## Supporting Information for "3D DNA structural barcode copying and random access"

### 1. 3D bit and DNA dumbbell unit design

The barcode '111111' are drawn in Figure S1. where the distance between two neighboring 3D bits is always 1032 bp. Barcodes '111', '011', and '001' are based on the same core barcode '111111' design. Barcodes '111', '011', and '001' have the first three, two or one 3D bit, respectively at the same position as in the core design.

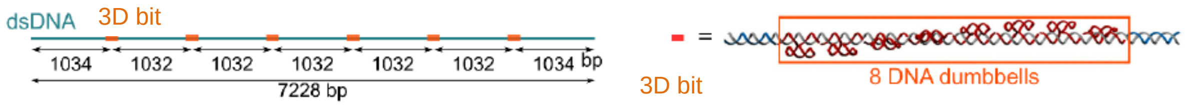

**Supplementary Figure S1.** The core design of barcode '111111' with the depicted structure of individual 3D bits that are equally interspaced (1032 bp) along the barcode.

Each 3D bit consists of eight DNA dumbbells. A DNA dumbbell oligonucleotide sequence has 48 nt and has three functional parts (Figure S2). First, 10 nt are complementary to the specific position in DNA scaffold, next 28 nt form DNA dumbbell with two four thymidine loops, and the last 10 nt are also complementary to DNA scaffold (Figure S2).

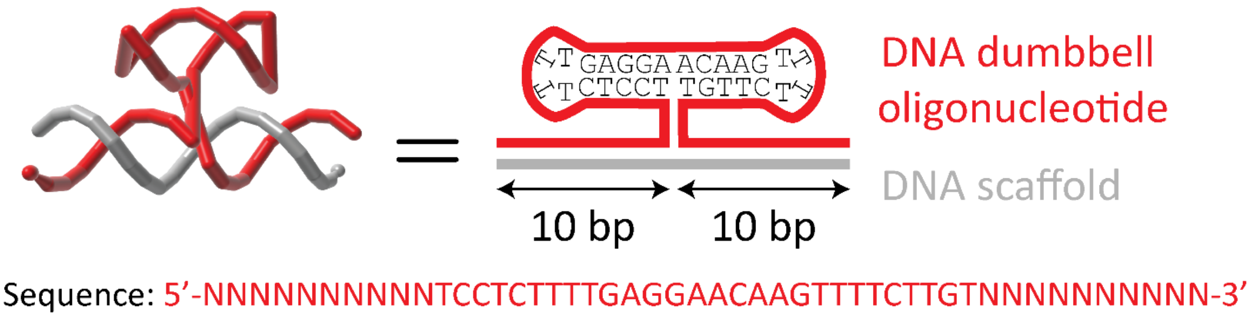

**Supplementary Figure S2.** DNA dumbbell oligonucleotide 3D and 2D sketch and sequence.

### 2. Oligonucleotide sequences

All DNA oligonucleotides used in this study were ordered from Integrated DNA Technologies (IDT) in IDTE buffer (10 mM Tris, 0.1 mM EDTA), 100 µM concentration and purified with standard desalting.

Oligonucleotides complementary to DNA scaffold (M13mp18) are shown in Table S1. From these 190 oligonucleotides, 188 have a length of 38 nt and terminal oligonucleotides have 46 nt in length and contain four thymidine nucleotides to minimalize potential a 3D DNA structural barcode aggregation.[1]

**Supplementary Table S1**. DNA scaffold complementary oligonucleotides

| Oligo number | Sequence (5’→3’) | Oligo Number | Sequence (5’→3’) |
|---|---|---|---|
| 1 | TTTTCGTAATCATGGTCATAGCTGTTTCCTGTGTGAAATTGTTATC | 96 | CTTGAGCCATTTGGGAATTAGAGCCAGCAAAATCACCA |
| 2 | CGCTCACAATTCCACACAACATACGAGCCGGAAGCATA | 97 | GTAGCACCATTACCATTAGCAAGGCCGGAAACGTCACC |
| 3 | AAGTGTAAAGCCTGGGGTGCCTAATGAGTGAGCTAACT | 98 | AATGAAACCATCGATAGCAGCACCGTAATCAGTAGCGA |
| 4 | CACATTAATTGCGTTGCGCTCACTGCCCGCTTTCCAGT | 99 | CAGAATCAAGTTTGCCTTTAGCGTCAGACTGTAGCGCG |
| 5 | CGGGAAACCTGTCGTGCCAGCTGCATTAATGAATCGGC | 100 | TTTTCATCGGCATTTTCGGTCATAGCCCCCTTATTAGC |
| 6 | CAACGCGCGGGGAGAGGCGGTTTGCGTATTGGGCGCCA | 101 | GTTTGCCATCTTTTCATAATCAAAATCACCGGAACCAG |
| 7 | GGGTGGTTTTTCTTTTCACCAGTGAGACGGGCAACAGC | 102 | AGCCACCACCGGAACCGCCTCCCTCAGAGCCGCCACCC |
| 8 | TGATTGCCCTTCACCGCCTGGCCCTGAGAGAGTTGCAG | 103 | TCAGAACCGCCACCCTCAGAGCCACCACCCTCAGAGCC |
| 9 | CAAGCGGTCCACGCTGGTTTGCCCCAGCAGGCGAAAAT | 104 | GCCACCAGAACCACCACCAGAGCCGCCGCCAGCATTGA |
| 10 | CCTGTTTGATGGTGGTTCCGAAATCGGCAAAATCCCTT | 105 | CAGGAGGTTGAGGCAGGTCAGACGATTGGCCTTGATAT |
| 11 | ATAAATCAAAAGAATAGCCCGAGATAGGGTTGAGTGTT | 106 | TCACAAACAAATAAATCCTCATTAAAGCCAGAATGGAA |
| 12 | GTTCCAGTTTGGAACAAGAGTCCACTATTAAAGAACGT | 107 | AGCGCAGTCTCTGAATTTACCGTTCCAGTAAGCGTCAT |
| 13 | GGACTCCAACGTCAAAGGGCGAAAAACCGTCTATCAGG | 108 | ACATGGCTTTTGATGATACAGGAGTGTACTGGTAATAA |
| 14 | GCGATGGCCCACTACGTGAACCATCACCCAAATCAAGT | 109 | GTTTTAACGGGGTCAGTGCCTTGAGTAACAGTGCCCGT |
| 15 | TTTTTGGGGTCGAGGTGCCGTAAAGCACTAAATCGGAA | 110 | ATAAACAGTTAATGCCCCCTGCCTATTTCGGAACCTAT |
| 16 | CCCTAAAGGGAGCCCCCGATTTAGAGCTTGACGGGGAA | 111 | TATTCTGAAACATGAAAGTATTAAGAGGCTGAGACTCC |
| 17 | AGCCGGCGAACGTGGCGAGAAAGGAAGGGAAGAAAGCG | 112 | TCAAGAGAAGGATTAGGATTAGCGGGGTTTTGCTCAGT |
| 18 | AAAGGAGCGGGCGCTAGGGCGCTGGCAAGTGTAGCGGT | 113 | ACCAGGCGGATAAGTGCCGTCGAGAGGGTTGATATAAG |
| 19 | CACGCTGCGCGTAACCACCACACCCGCCGCGCTTAATG | 114 | TATAGCCCGGAATAGGTGTATCACCGTACTCAGGAGGT |
| 20 | CGCCGCTACAGGGCGCGTACTATGGTTGCTTTGACGAG | 115 | TTAGTACCGCCACCCTCAGAACCGCCACCCTCAGAACC |
| 21 | CACGTATAACGTGCTTTCCTCGTTAGAATCAGAGCGGG | 116 | GCCACCCTCAGAGCCACCACCCTCATTTTCAGGGATAG |
| 22 | AGCTAAACAGGAGGCCGATTAAAGGGATTTTAGACAGG | 117 | CAAGCCCAATAGGAACCCATGTACCGTAACACTGAGTT |
| 23 | AACGGTACGCCAGAATCCTGAGAAGTGTTTTTATAATC | 118 | TCGTCACCAGTACAAACTACAACGCCTGTAGCATTCCA |
| 24 | AGTGAGGCCACCGAGTAAAAGAGTCTGTCCATCACGCA | 119 | CAGACAGCCCTCATAGTTAGCGTAACGATCTAAAGTTT |
| 25 | AATTAACCGTTGTAGCAATACTTCTTTGATTAGTAATA | 120 | TGTCGTCTTTCCAGACGTTAGTAAATGAATTTTCTGTA |
| 26 | ACATCACTTGCCTGAGTAGAAGAACTCAAACTATCGGC | 121 | TGGGATTTTGCTAAACAACTTTCAACAGTTTCAGCGGA |
| 27 | CTTGCTGGTAATATCCAGAACAATATTACCGCCAGCCA | 122 | GTGAGAATAGAAAGGAACAACTAAAGGAATTGCGAATA |
| 28 | TTGCAACAGGAAAAACGCTCATGGAAATACCTACATTT | 123 | ATAATTTTTTCACGTTGAAAATCTCCAAAAAAAAGGCT |
| 29 | TGACGCTCAATCGTCTGAAATGGATTATTTACATTGGC | 124 | CCAAAAGGAGCCTTTAATTGTATCGGTTTATCAGCTTG |
| 30 | AGATTCACCAGTCACACGACCAGTAATAAAAGGGACAT | 125 | CTTTCGAGGTGAATTTCTTAAACAGCTTGATACCGATA |
| 31 | TCTGGCCAACAGAGATAGAACCCTTCTGACCTGAAAGC | 126 | GTTGCGCCGACAATGACAACAACCATCGCCCACGCATA |
| 32 | GTAAGAATACGTGGCACAGACAATATTTTTGAATGGCT | 127 | ACCGATATATTCGGTCGCTGAGGCTTGCAGGGAGTTAA |
| 33 | ATTAGTCTTTAATGCGCGAACTGATAGCCCTAAAACAT | 128 | AGGCCGCTTTTGCGGGATCGTCACCCTCAGCAGCGAAA |
| 34 | CGCCATTAAAAATACCGAACGAACCACCAGCAGAAGAT | 129 | GACAGCATCGGAACGAGGGTAGCAACGGCTACAGAGGC |
| 35 | AAAACAGAGGTGAGGCGGTCAGTATTAACACCGCCTGC | 130 | TTTGAGGACTAAAGACTTTTTCATGAGGAAGTTTCCAT |
| 36 | AACAGTGCCACGCTGAGAGCCAGCAGCAAATGAAAAAT | 131 | TAAACGGGTAAAATACGTAATGCCACTACGAAGGCACC |
| 37 | CTAAAGCATCACCTTGCTGAACCTCAAATATCAAACCC | 132 | AACCTAAAACGAAAGAGGCAAAAGAATACACTAAAACA |
| 38 | TCAATCAATATCTGGTCAGTTGGCAAATCAACAGTTGA | 133 | CTCATCTTTGACCCCCAGCGATTATACCAAGCGCGAAA |
| 39 | AAGGAATTGAGGAAGGTTATCTAAAATATCTTTAGGAG | 134 | CAAAGTACAACGGAGATTTGTATCATCGCCTGATAAAT |
| 40 | CACTAACAACTAATAGATTAGAGCCGTCAATAGATAAT | 135 | TGTGTCGAAATCCGCGACCTGCTCCATGTTACTTAGCC |
| 41 | ACATTTGAGGATTTAGAAGTATTAGACTTTACAAACAA | 136 | GGAACGAGGCGCAGACGGTCAATCATAAGGGAACCGAA |
| 42 | TTCGACAACTCGTATTAAATCCTTTGCCCGAACGTTAT | 137 | CTGACCAACTTTGAAAGAGGACAGATGAACGGTGTACA |

| | | | |
|---|---|---|---|
| 43 | TAATTTTAAAAGTTTGAGTAACATTATCATTTTGCGGA | 138 | GACCAGGCGCATAGGCTGGCTGACCTTCATCAAGAGTA |
| 44 | ACAAAGAAACCACCAGAAGGAGCGGAATTATCATCATA | 139 | ATCTTGACAAGAACCGGATATTCATTACCCAAATCAAC |
| 45 | TTCCTGATTATCAGATGATGGCAATTCATCAATATAAT | 140 | GTAACAAAGCTGCTCATTCAGTGAATAAGGCTTGCCCT |
| 46 | CCTGATTGTTTGGATTATACTTCTGAATAATGGAAGGG | 141 | GACGAGAAACACCAGAACGAGTAGTAAATTGGGCTTGA |
| 47 | TTAGAACCTACCATATCAAAATTATTTGCACGTAAAAC | 142 | GATGGTTTAATTTCAACTTTAATCATTGTGAATTACCT |
| 48 | AGAAATAAAGAAATTGCGTAGATTTTCAGGTTTAACGT | 143 | TATGCGATTTTAAGAACTGGCTCATTATACCAGTCAGG |
| 49 | CAGATGAATATACAGTAACAGTACCTTTTACATCGGGA | 144 | ACGTTGGGAAGAAAAATCTACGTTAATAAAACGAACTA |
| 50 | GAAACAATAACGGATTCGCCTGATTGCTTTGAATACCA | 145 | ACGGAACAACATTATTACAGGTAGAAAGATTCATCAGT |
| 51 | AGTTACAAAATCGCGCAGAGGCGAATTATTCATTTCAA | 146 | TGAGATTTAGGAATACCACATTCAACTAATGCAGATAC |
| 52 | TTACCTGAGCAAAAGAAGATGATGAAACAAACATCAAG | 147 | ATAACGCCAAAAGGAATTACGAGGCATAGTAAGAGCAA |
| 53 | AAAACAAAATTAATTACATTTAACAATTTCATTTGAAT | 148 | CACTATCATAACCCTCGTTTACCAGACGACGATAAAAA |
| 54 | TACCTTTTTAATGGAAACAGTACATAAATCAATATAT | 149 | CCAAAATAGCGAGAGGCTTTTGCAAAAGAAGTTTTGCC |
| 55 | GTGAGTGAATAACCTTGCTTCTGTAAATCGTCGCTATT | 150 | AGAGGGGGTAATAGTAAAATGTTTAGACTGGATAGCGT |
| 56 | AATTAATTTTCCCTTAGAATCCTTGAAAACATAGCGAT | 151 | CCAATACTGCGGAATCGTCATAAATATTCATTGAATCC |
| 57 | AGCTTAGATTAAGACGCTGAGAAGAGTCAATAGTGAAT | 152 | CCCTCAAATGCTTTAAACAGTTCAGAAAACGAGAATGA |
| 58 | TTATCAAAATCATAGGTCTGAGAGACTACCTTTTTAAC | 153 | CCATAAATCAAAAATCAGGTCTTTACCCTGACTATTAT |
| 59 | CTCCGGCTTAGGTTGGGTTATATAACTATATGTAAATG | 154 | AGTCAGAAGCAAAGCGGATTGCATCAAAAAGATTAAGA |
| 60 | CTGATGCAAATCCAATCGCAAGACAAAGAACGCGAGAA | 155 | GGAAGCCCGAAAGACTTCAAATATCGCGTTTTAATTCG |
| 61 | AACTTTTTCAAATATATTTTAGTTAATTTCATCTTCTG | 156 | AGCTTCAAAGCGAACCAGACCGGAAGCAAACTCCAACA |
| 62 | ACCTAAATTTAATGGTTTGAAATACCGACCGTGTGATA | 157 | GGTCAGGATTAGAGAGTACCTTTAATTGCTCCTTTTGA |
| 63 | AATAAGGCGTTAAATAAGAATAAACACCGGAATCATAA | 158 | TAAGAGGTCATTTTTGCGGATGGCTTAGAGCTTAATTG |
| 64 | TTACTAGAAAAAGCCTGTTTAGTATCATATGCGTTATA | 159 | CTGAATATAATGCTGTAGCTCAACATGTTTTAAATATG |
| 65 | CAAATTCTTACCAGTATAAAGCCAACGCTCAACAGTAG | 160 | CAACTAAAGTACGGTGTCTGGAAGTTTCATTCCATATA |
| 66 | GGCTTAATTGAGAATCGCCATATTTAACAACGCCAACA | 161 | ACAGTTGATTCCCAATTCTGCGAACGAGTAGATTTAGT |
| 67 | TGTAATTTAGGCAGAGGCATTTTCGAGCCAGTAATAAG | 162 | TTGACCATTAGATACATTTCGCAAATGGTCAATAACCT |
| 68 | AGAATATAAAGTACCGACAAAAGGTAAAGTAATTCTGT | 163 | GTTTAGCTATATTTTCATTTGGGGCGCGAGCTGAAAAG |
| 69 | CCAGACGACGACAATAAACAACATGTTCAGCTAATGCA | 164 | GTGGCATCAATTCTACTAATAGTAGTAGCATTAACATC |
| 70 | GAACGCGCCTGTTTATCAACAATAGATAAGTCCTGAAC | 165 | CAATAAATCATACAGGCAAGGCAAAGAATTAGCAAAAT |
| 71 | AAGAAAAATAATATCCCATCCTAATTTACGAGCATGTA | 166 | TAAGCAATAAAGCCTCAGAGCATAAAGCTAAATCGGTT |
| 72 | GAAACCAATCAATAATCGGCTGTCTTTCCTTATCATTC | 167 | GTACCAAAAACATTATGACCCTGTAATACTTTTGCGGG |
| 73 | CAAGAACGGGTATTAAACCAAGTACCGCACTCATCGAG | 168 | AGAAGCCTTTATTTCAACGCAAGGATAAAAATTTTTAG |
| 74 | AACAAGCAAGCCGTTTTTATTTTCATCGTAGGAATCAT | 169 | AACCCTCATATATTTTAAATGCAATGCCTGAGTAATGT |
| 75 | TACCGCGCCCAATAGCAAGCAAATCAGATATAGAAGGC | 170 | GTAGGTAAAGATTCAAAAGGGTGAGAAAGGCCGGAGAC |
| 76 | TTATCCGGTATTCTAAGAACGCGAGGCGTTTTAGCGAA | 171 | AGTCAAATCACCATCAATATGATATTCAACCGTTCTAG |
| 77 | CCTCCCGACTTGCGGGAGGTTTTGAAGCCTTAAATCAA | 172 | CTGATAAATTAATGCCGGAGAGGGTAGCTATTTTTGAG |
| 78 | GATTAGTTGCTATTTTGCACCCAGCTACAATTTTATCC | 173 | AGATCTACAAAGGCTATCAGGTCATTGCCTGAGAGTCT |
| 79 | TGAATCTTACCAACGCTAACGAGCGTCTTTCCAGAGCC | 174 | GGAGCAAACAAGAGAATCGATGAACGGTAATCGTAAAA |
| 80 | TAATTTGCCAGTTACAAAATAAACAGCCATATTATTTA | 175 | CTAGCATGTCAATCATATGTACCCCGGTTGATAATCAG |
| 81 | TCCCAATCCAAATAAGAAACGATTTTTTGTTTAACGTC | 176 | AAAAGCCCCAAAAACAGGAAGATTGTATAAGCAAATAT |
| 82 | AAAAATGAAAATAGCAGCCTTTACAGAGAGAATAACAT | 177 | TTAAATTGTAAACGTTAATATTTTGTTAAAATTCGCAT |
| 83 | AAAAACAGGGAAGCGCATTAGACGGGAGAATTAACTGA | 178 | TAAATTTTTGTTAAATCAGCTCATTTTTTAACCAATAG |
| 84 | ACACCCTGAACAAAGTCAGAGGGTAATTGAGCGCTAAT | 179 | GAACGCCATCAAAAATAATTCGCGTCTGGCCTTCCTGT |
| 85 | ATCAGAGAGATAACCCACAAGAATTGAGTTAAGCCCAA | 180 | AGCCAGCTTTCATCAACATTAAATGTGAGCGAGTAACA |
| 86 | TAATAAGAGCAAGAAACAATGAAATAGCAATAGCTATC | 181 | ACCCGTCGGATTCTCCGTGGGAACAAACGGCGGATTGA |
| 87 | TTACCGAAGCCCTTTTTAAGAAAAGTAAGCAGATAGCC | 182 | CCGTAATGGGATAGGTCACGTTGGTGTAGATGGGCGCA |
| 88 | GAACAAAGTTACCAGAAGGAAACCGAGGAAACGCAATA | 183 | TCGTAACCGTGCATCTGCCAGTTTGAGGGGACGACGAC |
| 89 | ATAACGGAATACCCAAAAGAACTGGCATGATTAAGACT | 184 | AGTATCGGCCTCAGGAAGATCGCACTCCAGCCAGCTTT |
| 90 | CCTTATTACGCAGTATGTTAGCAAACGTAGAAAATACA | 185 | CCGGCACCGCTTCTGGTGCCGGAAACCAGGCAAAGCGC |
| 91 | TACATAAAGGTGGCAACATATAAAAGAAACGCAAAGAC | 186 | CATTCGCCATTCAGGCTGCGCAACTGTTGGGAAGGGCG |
| 92 | ACCACGGAATAAGTTTATTTTGTCACAATCAATAGAAA | 187 | ATCGGTGCGGGCCTCTTCGCTATTACGCCAGCTGGCGA |

| | | | |
|---|---|---|---|
| 93 | ATTCATATGGTTTACCAGCGCCAAAGACAAAAGGGCGA | 188 | AAGGGGGATGTGCTGCAAGGCGATTAAGTTGGGTAACG |
| 94 | CATTCAACCGATTGAGGGAGGGAAGGTAAATATTGACG | 189 | CCAGGGTTTTCCCAGTCACGACGTTGTAAAACGACGGC |
| 95 | GAAATTATTCATTAAAGGTGAATTATCACCGTCACCGA | 190 | CAGTGCCAAGCTTGCATGCCTGCAGGTCGACTCTAGAGGATCT[illegible] |

To assemble a 3D DNA structural barcode some of the oligonucleotides form Table S1 were replaced with oligonucleotides that contain the DNA dumbbell sequence (Figure S1). Numbers of oligonucleotides that were replaced in the mixture of 190 oligonucleotides shown in Table S1 are listed for a specific barcode in Table S2, S3, S4, and S5 and correspond to the designed 3D DNA structural barcodes '111111', '111', '011', and '001' respectively.

**Supplementary Table S2.** Barcode '111111' replaced oligonucleotides with the provided sequences.

| 3D bit number | Oligonucleotide number | Sequence (5'→'3') | Length (nt) | To replace oligonucleotides |
|---|---|---|---|---|
| 1 | A1 | ACATCACTTGTCCTCTTTTGAGGAACAAGTTTTCTTGTCCTGAGTAGA | 48 | 26-30 |
| | A2 | AGAACTCAAATCCTCTTTTGAGGAACAAGTTTTCTTGTCTATCGGCCT | 48 | |
| | A3 | TGCTGGTAATTCCTCTTTTGAGGAACAAGTTTTCTTGTATCCAGAACA | 48 | |
| | A4 | ATATTACCGCTCCTCTTTTGAGGAACAAGTTTTCTTGTCAGCCATTGC | 48 | |
| | A5 | AACAGGAAAATCCTCTTTTGAGGAACAAGTTTTCTTGTACGCTCATGG | 48 | |
| | A6 | AAATACCTACTCCTCTTTTGAGGAACAAGTTTTCTTGTATTTTGACGC | 48 | |
| | A7 | TCAATCGTCTTCCTCTTTTGAGGAACAAGTTTTCTTGTGAAATGGATT | 48 | |
| | A8 | ATTTACATTGTCCTCTTTTGAGGAACAAGTTTTCTTGTGCAGATTCAC | 48 | |
| | A9 | CAGTCACACGACCAGTAATAAAAGGGACAT | 30 | |
| 2 | B1 | TTACCTGAGCAAAAGAAGATGATGAAACAAACATCAAGAAAACA | 44 | 52-57 |
| | B2 | AAATTAATTATCCTCTTTTGAGGAACAAGTTTTCTTGTCATTTAACAA | 48 | |
| | B3 | TTTCATTTGATCCTCTTTTGAGGAACAAGTTTTCTTGTATTACCTTTT | 48 | |
| | B4 | TTAATGGAAATCCTCTTTTGAGGAACAAGTTTTCTTGTCAGTACATAA | 48 | |
| | B5 | ATCAATATATTCCTCTTTTGAGGAACAAGTTTTCTTGTGTGAGTGAAT | 48 | |
| | B6 | AACCTTGCTTTCCTCTTTTGAGGAACAAGTTTTCTTGTCTGTAAATCG | 48 | |
| | B7 | TCGCTATTAATCCTCTTTTGAGGAACAAGTTTTCTTGTTTAATTTTCC | 48 | |
| | B8 | CTTAGAATCCTCCTCTTTTGAGGAACAAGTTTTCTTGTTTGAAAACAT | 48 | |
| | B9 | AGCGATAGCTTCCTCTTTTGAGGAACAAGTTTTCTTGTTAGATTAAGA | 48 | |
| | B10 | CGCTGAGAAGAGTCAATAGTGAAT | 24 | |
| 3 | C1 | TGAATCTTACCAACGCTAACGAGCGTCTTTCCAGAGCCTAATTTGCCAGT | 50 | 79-85 |
| | C2 | TACAAAATAATCCTCTTTTGAGGAACAAGTTTTCTTGTACAGCCATAT | 48 | |
| | C3 | TATTTATCCCTCCTCTTTTGAGGAACAAGTTTTCTTGTAATCCAAATA | 48 | |
| | C4 | AGAAACGATTTCCTCTTTTGAGGAACAAGTTTTCTTGTTTTTGTTTAA | 48 | |
| | C5 | CGTCAAAAAATTCCTCTTTTGAGGAACAAGTTTTCTTGTGAAAATAGCA | 48 | |
| | C6 | GCCTTTACAGTCCTCTTTTGAGGAACAAGTTTTCTTGTAGAGAATAAC | 48 | |
| | C7 | ATAAAAACAGTCCTCTTTTGAGGAACAAGTTTTCTTGTGGAAGCGCAT | 48 | |
| | C8 | TAGACGGGAGTCCTCTTTTGAGGAACAAGTTTTCTTGTAATTAACTGA | 48 | |
| | C9 | ACACCCTGAATCCTCTTTTGAGGAACAAGTTTTCTTGTCAAAGTCAGA | 48 | |
| | C10 | GGGTAATTGAGCGCTAATATCAGAGAGATAACCCACAAGAATTGAGTTAAGCCCAA | 56 | |
| 4 | D1 | TCACAAACAAATAAATCCTCATTAAAGCCAGAATGGAAAGCGCAGTCTCTGAATTT | 56 | 106-112 |
| | D2 | ACCGTTCCAGTCCTCTTTTGAGGAACAAGTTTTCTTGTTAAGCGTCAT | 48 | |
| | D3 | ACATGGCTTTTCCTCTTTTGAGGAACAAGTTTTCTTGTTGATGATACA | 48 | |
| | D4 | GGAGTGTACTTCCTCTTTTGAGGAACAAGTTTTCTTGTGGTAATAAGT | 48 | |

| | | | | |
|---|---|---|---|---|
| | D5 | TTTAACGGGGTCCTCTTTTGAGGAACAAGTTTTCTTGTTCAGTGCCTT | 48 | |
| | D6 | GAGTAACAGTTCCTCTTTTGAGGAACAAGTTTTCTTGTGCCCGTATAA | 48 | |
| | D7 | ACAGTTAATGTCCTCTTTTGAGGAACAAGTTTTCTTGTCCCCCTGCCT | 48 | |
| | D8 | ATTTCGGAACTCCTCTTTTGAGGAACAAGTTTTCTTGTCTATTATTCT | 48 | |
| | D9 | GAAACATGAATCCTCTTTTGAGGAACAAGTTTTCTTGTAGTATTAAGA | 48 | |
| | D10 | GGCTGAGACTCCTCAAGAGAAGGATTAGGATTAGCGGGGTTTTGCTCAGT | 50 | |
| 5 | E1 | CAAAGTACAACGGAGATTTGTATC | 24 | 134-139 |
| | E2 | ATCGCCTGATTCCTCTTTTGAGGAACAAGTTTTCTTGTAAATTGTGTC | 48 | |
| | E3 | GAAATCCGCGTCCTCTTTTGAGGAACAAGTTTTCTTGTACCTGCTCCA | 48 | |
| | E4 | TGTTACTTAGTCCTCTTTTGAGGAACAAGTTTTCTTGTCCGGAACGAG | 48 | |
| | E5 | GCGCAGACGGTCCTCTTTTGAGGAACAAGTTTTCTTGTTCAATCATAA | 48 | |
| | E6 | GGGAACCGAATCCTCTTTTGAGGAACAAGTTTTCTTGTCTGACCAACT | 48 | |
| | E7 | TTGAAAGAGGTCCTCTTTTGAGGAACAAGTTTTCTTGTACAGATGAAC | 48 | |
| | E8 | GGTGTACAGATCCTCTTTTGAGGAACAAGTTTTCTTGTCCAGGCGCAT | 48 | |
| | E9 | AGGCTGGCTGTCCTCTTTTGAGGAACAAGTTTTCTTGTACCTTCATCA | 48 | |
| | E10 | AGAGTAATCTTGACAAGAACCGGATATTCATTACCCAAATCAAC | 44 | |
| 6 | F1 | ACAGTTGATTCCCAATTCTGCGAACGAGTA | 30 | 161-165 |
| | F2 | GATTTAGTTTTCCTCTTTTGAGGAACAAGTTTTCTTGTGACCATTAGA | 48 | |
| | F3 | TACATTTCGCTCCTCTTTTGAGGAACAAGTTTTCTTGTAAATGGTCAA | 48 | |
| | F4 | TAACCTGTTTTCCTCTTTTGAGGAACAAGTTTTCTTGTAGCTATATTT | 48 | |
| | F5 | TCATTTGGGGTCCTCTTTTGAGGAACAAGTTTTCTTGTCGCGAGCTGA | 48 | |
| | F6 | AAAGGTGGCATCCTCTTTTGAGGAACAAGTTTTCTTGTTCAATTCTAC | 48 | |
| | F7 | TAATAGTAGTTCCTCTTTTGAGGAACAAGTTTTCTTGTAGCATTAACA | 48 | |
| | F8 | TCCAATAAATTCCTCTTTTGAGGAACAAGTTTTCTTGTCATACAGGCA | 48 | |
| | F9 | AGGCAAAGAATCCTCTTTTGAGGAACAAGTTTTCTTGTTTAGCAAAAT | 48 | |

**Supplementary Table S3.** Barcode '111' replaced oligonucleotides with the provided sequences.

| 3D bit number | Oligo number | Sequence (5'→3') | Length (nt) | To replace oligos |
|---|---|---|---|---|
| 1 | A1 | ACATCACTTGTCCTCTTTTGAGGAACAAGTTTTCTTGTCCTGAGTAGA | 48 | 26-30 |
| | A2 | AGAACTCAAATCCTCTTTTGAGGAACAAGTTTTCTTGTCTATCGGCCT | 48 | |
| | A3 | TGCTGGTAATTCCTCTTTTGAGGAACAAGTTTTCTTGTATCCAGAACA | 48 | |
| | A4 | ATATTACCGCTCCTCTTTTGAGGAACAAGTTTTCTTGTCAGCCATTGC | 48 | |
| | A5 | AACAGGAAAATCCTCTTTTGAGGAACAAGTTTTCTTGTACGCTCATGG | 48 | |
| | A6 | AAATACCTACTCCTCTTTTGAGGAACAAGTTTTCTTGTATTTTGACGC | 48 | |
| | A7 | TCAATCGTCTTCCTCTTTTGAGGAACAAGTTTTCTTGTGAAATGGATT | 48 | |
| | A8 | ATTTACATTGTCCTCTTTTGAGGAACAAGTTTTCTTGTGCAGATTCAC | 48 | |
| | A9 | CAGTCACACGACCAGTAATAAAAGGGACAT | 30 | |
| 2 | B1 | TTACCTGAGCAAAAGAAGATGATGAAACAAACATCAAGAAAACA | 44 | 52-57 |
| | B2 | AAATTAATTATCCTCTTTTGAGGAACAAGTTTTCTTGTCATTTAACAA | 48 | |
| | B3 | TTTCATTTGATCCTCTTTTGAGGAACAAGTTTTCTTGTATTACCTTTT | 48 | |
| | B4 | TTAATGGAAATCCTCTTTTGAGGAACAAGTTTTCTTGTCAGTACATAA | 48 | |
| | B5 | ATCAATATATTCCTCTTTTGAGGAACAAGTTTTCTTGTGTGAGTGAAT | 48 | |
| | B6 | AACCTTGCTTTCCTCTTTTGAGGAACAAGTTTTCTTGTCTGTAAATCG | 48 | |
| | B7 | TCGCTATTAATCCTCTTTTGAGGAACAAGTTTTCTTGTTTAATTTTCC | 48 | |
| | B8 | CTTAGAATCCTCCTCTTTTGAGGAACAAGTTTTCTTGTTTGAAAACAT | 48 | |
| | B9 | AGCGATAGCTTCCTCTTTTGAGGAACAAGTTTTCTTGTTAGATTAAGA | 48 | |
| | B10 | CGCTGAGAAGAGTCAATAGTGAAT | 24 | |
| 3 | C1 | TGAATCTTACCAACGCTAACGAGCGTCTTTCCAGAGCCTAATTTGCCAGT | 50 | 79-85 |
| | C2 | TACAAAATAATCCTCTTTTGAGGAACAAGTTTTCTTGTACAGCCATAT | 48 | |
| | C3 | TATTTATCCCTCCTCTTTTGAGGAACAAGTTTTCTTGTAATCCAAATA | 48 | |
| | C4 | AGAAACGATTTCCTCTTTTGAGGAACAAGTTTTCTTGTTTTTGTTTAA | 48 | |
| | C5 | CGTCAAAAAATTCCTCTTTTGAGGAACAAGTTTTCTTGTGAAAATAGCA | 48 | |
| | C6 | GCCTTTACAGTCCTCTTTTGAGGAACAAGTTTTCTTGTAGAGAATAAC | 48 | |
| | C7 | ATAAAAACAGTCCTCTTTTGAGGAACAAGTTTTCTTGTGGAAGCGCAT | 48 | |
| | C8 | TAGACGGGAGTCCTCTTTTGAGGAACAAGTTTTCTTGTAATTAACTGA | 48 | |
| | C9 | ACACCCTGAATCCTCTTTTGAGGAACAAGTTTTCTTGTCAAAGTCAGA | 48 | |
| | C10 | GGGTAATTGAGCGCTAATATCAGAGAGATAACCCACAAGAATTGAGTTAAGCCCAA | 56 | |

**Supplementary Table S4.** Barcode '011' replaced oligonucleotides with the provided sequences.

| 3D bit number | Oligo number | Sequence (5'→3') | Length (nt) | To replace oligos |
|---|---|---|---|---|
| 2 | B1 | TTACCTGAGCAAAAGAAGATGATGAAACAAACATCAAGAAAACA | 44 | 52-57 |
| | B2 | AAATTAATTATCCTCTTTTGAGGAACAAGTTTTCTTGTCATTTAACAA | 48 | |
| | B3 | TTTCATTTGATCCTCTTTTGAGGAACAAGTTTTCTTGTATTACCTTTT | 48 | |
| | B4 | TTAATGGAAATCCTCTTTTGAGGAACAAGTTTTCTTGTCAGTACATAA | 48 | |
| | B5 | ATCAATATATTCCTCTTTTGAGGAACAAGTTTTCTTGTGTGAGTGAAT | 48 | |
| | B6 | AACCTTGCTTTCCTCTTTTGAGGAACAAGTTTTCTTGTCTGTAAATCG | 48 | |
| | B7 | TCGCTATTAATCCTCTTTTGAGGAACAAGTTTTCTTGTTTAATTTTCC | 48 | |
| | B8 | CTTAGAATCCTCCTCTTTTGAGGAACAAGTTTTCTTGTTTGAAAACAT | 48 | |
| | B9 | AGCGATAGCTTCCTCTTTTGAGGAACAAGTTTTCTTGTTAGATTAAGA | 48 | |
| | B10 | CGCTGAGAAGAGTCAATAGTGAAT | 24 | |
| 3 | C1 | TGAATCTTACCAACGCTAACGAGCGTCTTTCCAGAGCCTAATTTGCCAGT | 50 | 79-85 |
| | C2 | TACAAAATAATCCTCTTTTGAGGAACAAGTTTTCTTGTACAGCCATAT | 48 | |

| | | |
|---|---|---|
| C3 | TATTTATCCCTCCTCTTTTGAGGAACAAGTTTTCTTGTAATCCAAATA | 48 |
| C4 | AGAAACGATTTCCTCTTTTGAGGAACAAGTTTTCTTGTTTTTGTTTAA | 48 |
| C5 | CGTCAAAAATTCCTCTTTTGAGGAACAAGTTTTCTTGTGAAAATAGCA | 48 |
| C6 | GCCTTTACAGTCCTCTTTTGAGGAACAAGTTTTCTTGTAGAGAATAAC | 48 |
| C7 | ATAAAAACAGTCCTCTTTTGAGGAACAAGTTTTCTTGTGGAAGCGCAT | 48 |
| C8 | TAGACGGGAGTCCTCTTTTGAGGAACAAGTTTTCTTGTAATTAACTGA | 48 |
| C9 | ACACCCTGAATCCTCTTTTGAGGAACAAGTTTTCTTGTCAAAGTCAGA | 48 |
| C10 | GGGTAATTGAGCGCTAATATCAGAGAGATAACCCACAAGAATTGAGTTAAGCCCAA | 56 |

**Supplementary Table S5.** Barcode ‘001’ replaced oligonucleotides with the provided sequences.

| 3D bit number | Oligo number | Sequence (5’→’3’) | Length (nt) | To replace oligos |
|---|---|---|---|---|
| 3 | C1 | TGAATCTTACCAACGCTAACGAGCGTCTTTCCAGAGCCTAATTTGCCAGT | 50 | 79-85 |
| | C2 | TACAAAATAATCCTCTTTTGAGGAACAAGTTTTCTTGTACAGCCATAT | 48 | |
| | C3 | TATTTATCCCTCCTCTTTTGAGGAACAAGTTTTCTTGTAATCCAAATA | 48 | |
| | C4 | AGAAACGATTTCCTCTTTTGAGGAACAAGTTTTCTTGTTTTTGTTTAA | 48 | |
| | C5 | CGTCAAAAATTCCTCTTTTGAGGAACAAGTTTTCTTGTGAAAATAGCA | 48 | |
| | C6 | GCCTTTACAGTCCTCTTTTGAGGAACAAGTTTTCTTGTAGAGAATAAC | 48 | |
| | C7 | ATAAAAACAGTCCTCTTTTGAGGAACAAGTTTTCTTGTGGAAGCGCAT | 48 | |
| | C8 | TAGACGGGAGTCCTCTTTTGAGGAACAAGTTTTCTTGTAATTAACTGA | 48 | |
| | C9 | ACACCCTGAATCCTCTTTTGAGGAACAAGTTTTCTTGTCAAAGTCAGA | 48 | |
| | C10 | GGGTAATTGAGCGCTAATATCAGAGAGATAACCCACAAGAATTGAGTTAAGCCCAA | 56 | |

### 3. Nanopore data analysis workflow

From current data traces, we isolated single 3D DNA structural barcode events (Figure S3A) based on the current threshold ($\Delta I_{threshold}$) measured from the base current ($I_0$). After isolation of individual events, folded barcode events are discarded (Figure S3B) based on the current threshold set for the beginning of the event. In further analysis only unfolded barcode events were used (Figure S3C) and barcode readout is determined based on the number and normalized position to the beginning of 3D bits.

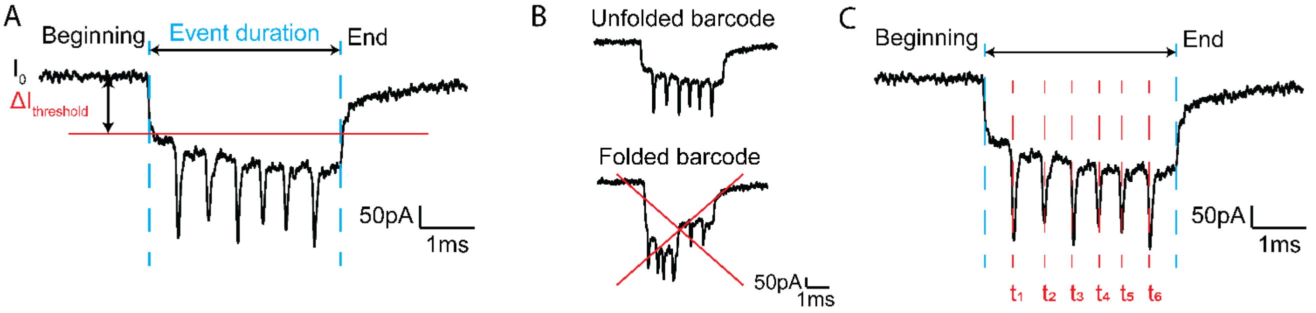

**Supplementary Figure S3.** Steps in nanopore data analysis of a 3D DNA structural barcode events. An Isolation of individual barcode events based on current drop threshold ($\Delta I_{threshold}$), event duration ($\Delta t$) and event charge deficit

(ECD). B The next step includes separation of unfolded form folded barcode events. C Only unfolded barcode events were used in further analysis of barcodes number based on normalized peak position of each 3D unit ($t_{1\text{-}6}$).

Data were analyzed using custom made LabView codes. Further details of data analysis of nanopore measurements can be found in previous group research. (1)

### 4. PCR primers and barcode-specific terminal sequences

Terminal sequences of barcodes shown in Table S6. are replaced with barcode-specific terminal sequences that serve as a template for the random-access of the barcode as presented in Figure 4. Highlighted in different colors are sequence part that is non-complementary to DNA scaffold ends.

**Supplementary Table S6.** Terminal sequences specific to barcodes '111', '011', and '001'.

| Barcode | Oligo Number | Sequence (5'→'3') | Length (nt) | To replace oligos |
|---|---|---|---|---|
| '111' | 111_beginning | TTTTGGTCTTGACAAACGTGTGCTTTTCCTGTGTGAAATTGTTATC | 46 | 1 |
| | 111_end | CAGTGCCAAGCTTGCATGCCTGCCGATGTTGACGGACTAATCTTTT | 46 | 190 |
| '011' | 011_beginning | TTTTTATCCCGTGAAGCTTGAGTGTTTCCTGTGTGAAATTGTTATC | 46 | 1 |
| | 011_end | CAGTGCCAAGCTTGCATGCCTGGGTATGGCACGCCTAATCTGTTTT | 46 | 190 |
| '001' | 001_beginning | TTTTTATGAGGACGAATCTCCCGCTTTCCTGTGTGAAATTGTTATC | 46 | 1 |
| | 001_end | CAGTGCCAAGCTTGCATGCCTGCCGTACCTAGATACACTCAATTTT | 46 | 190 |

Primers (Table S7) were ordered from IDT in nuclease-free water, 100 µM concentration and purified with standard desalting.

**Supplementary Table S7.** PCR primers used in this study.

| Primer name | Sequence (5'→'3') | Length (nt) |
|---|---|---|
| M13 forward primer | TTTTCGTAATCATGGTCATAGCTG | 24 |
| M13 reverse primer | AAAAGATCCTCTAGAGTCGACCTG | 24 |
| Barcode '111' primer | AAAAGATTAGTCCGTCAACATCGG | 24 |

### 5. Example events

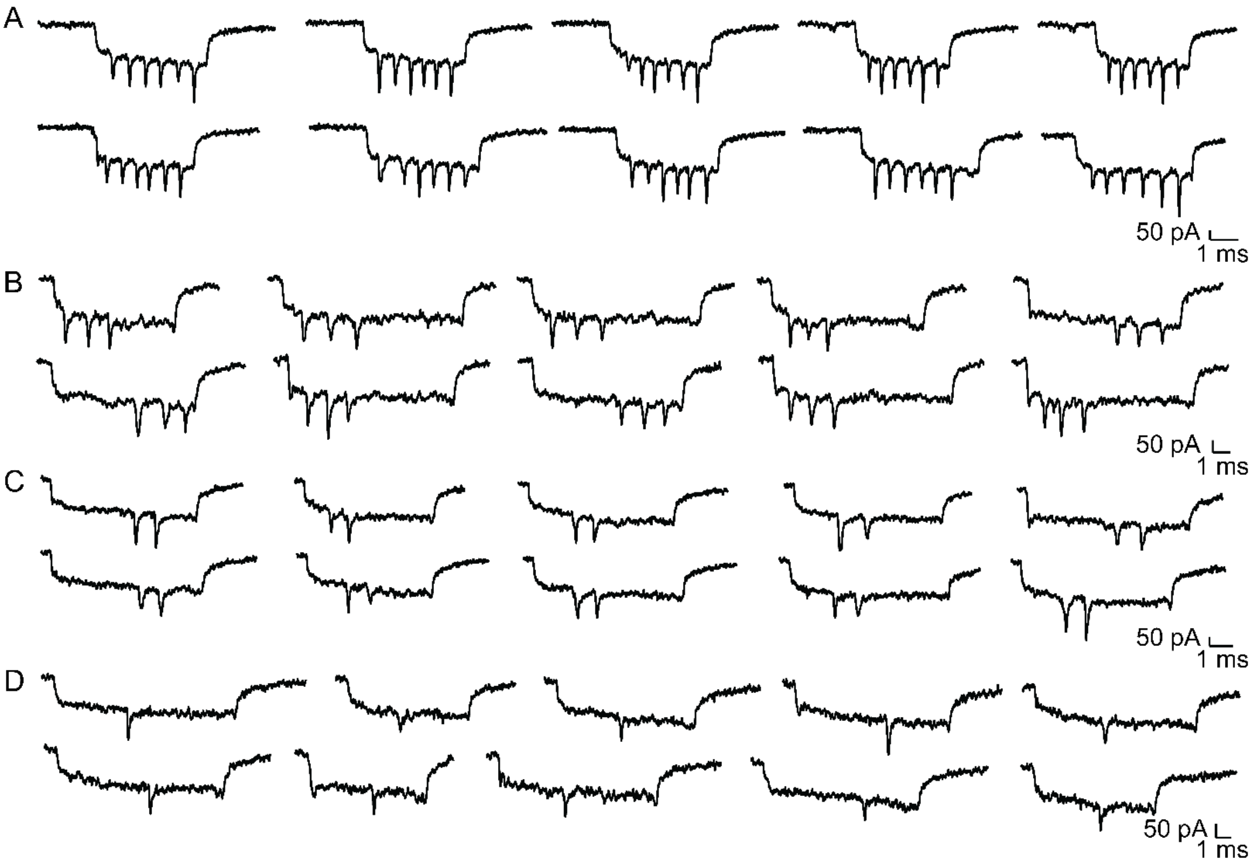

**Supplementary Figure S4.** Example events of nanopore readout barcodes shown in Figure 2. before PCR amplification. Events for barcode '111111', '111', '011', and '001' are shown in A, B, C and D, respectively.

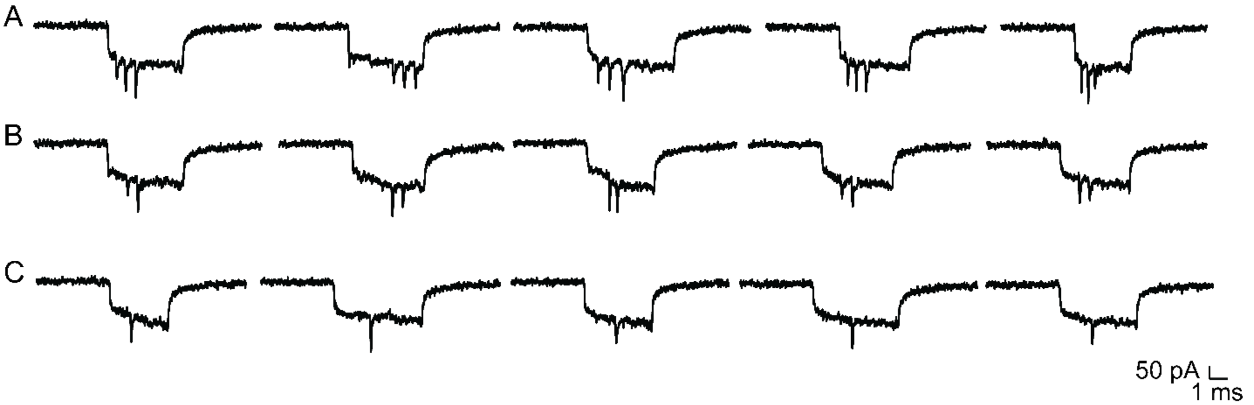

**Supplementary Figure S5.** Example events from nanopore readout of an equimolar mixture of barcodes '111', '011', and '001' are shown as A, B, and C, respectively. Example events are for analysis shown Figure 4. before PCR amplification.

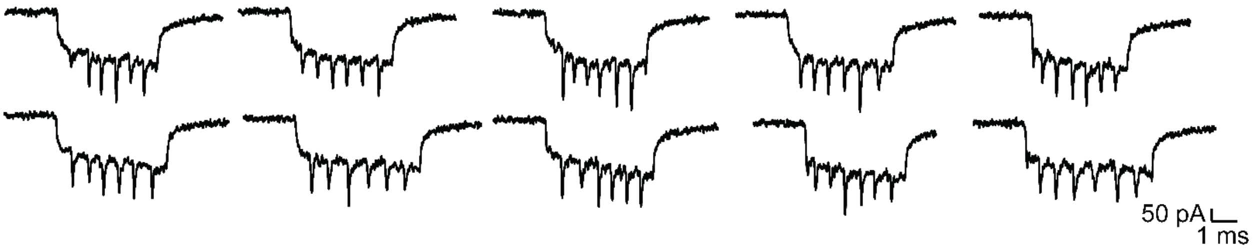

**Supplementary Figure S6.** Example events from nanopore readout of barcodes shown in Figure 3. after PCR amplification.

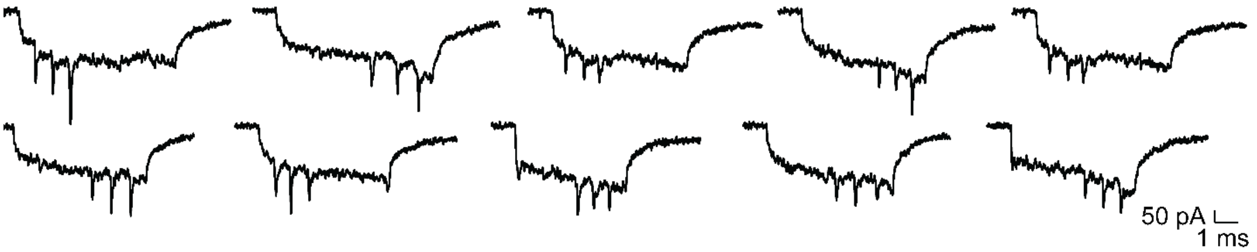

**Supplementary Figure S7.** Example events from nanopore readout of barcodes shown in Figure 4. after PCR amplification.

### 6. 3D DNA structural barcodes characterization

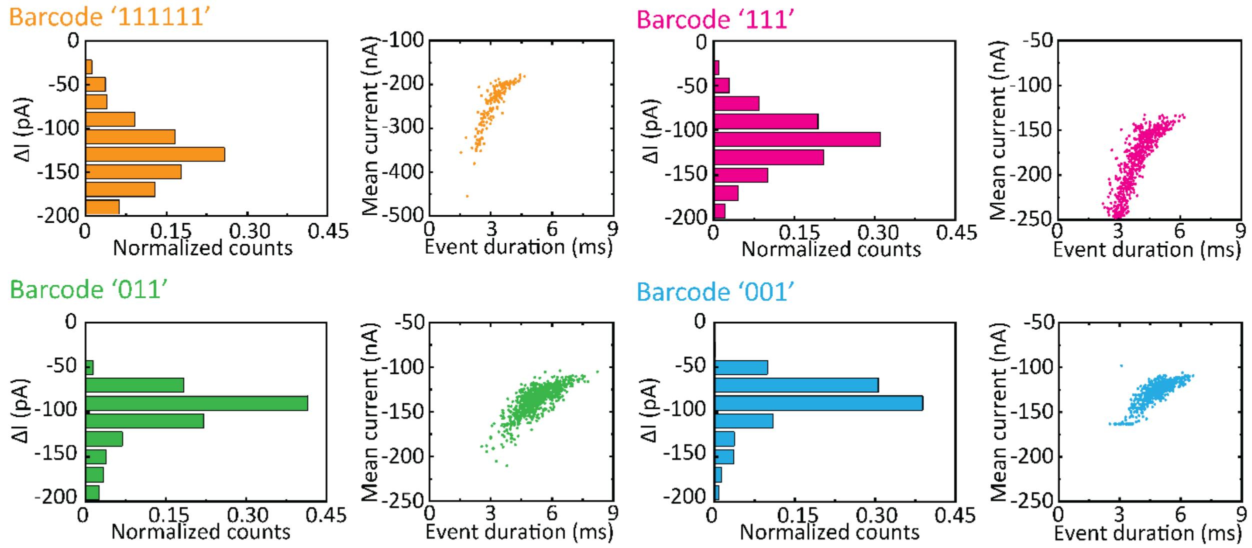

**Supplementary Figure S8.** 3D DNA structural barcodes additional features from Figure 2. Normalized histogram of 3D bit current drop is shown for all detected 3D bits (ΔI). Scatter plot of mean current (ECD divided by event duration) versus event duration of individual barcode events.
